# Obstacle Negotiation in Female Locust Oviposition Digging

**DOI:** 10.1101/2024.03.05.583468

**Authors:** Chen Klechevski, Lazar Kats, Amir Ayali

## Abstract

The female locust lays its eggs deep within soft substrate to protect them from predators and provide optimal conditions for successful hatching. During oviposition digging, the female’s abdomen extends into the ground, guided by a dedicated excavation mechanism at its tip, comprising two pairs of specialized digging valves. Little is known about how these active valves negotiate the various obstacles encountered on their path. In this study, female locusts oviposited their eggs in specialized sand-filled tubes with pre-inserted 3D-printed plastic obstacles. The subterranean route taken by the abdomen and digging valves upon encountering the obstacles was investigated, characterized, and compared to that in control tubes without obstacles. Data were obtained by way of visual inspection, by utilizing Cone Beam Computed Tomography scans in high-definition mode, and by making paraffin casts of the oviposition burrows (after egg hatching). We demonstrate, for the first time, the subterranean navigation ability of the female locust’s excavation mechanism and its ability to circumvent obstacles during oviposition. Finally, we discuss the role of active sensory-motor mechanisms versus the passive embodied function of the valves, central control, and decision-making.

## Introduction

Many insects use elongated ovipositors – the specialized apparatus at the tip of the female insect abdomen – to deposit their eggs in a suitable, often difficult to reach, moist and safe environment (e.g., Emeljanov, 2014; Elias et al., 2018; Cerkvenik et al., 2019). The female of the desert locust lacks such long appendages, and instead extends its abdomen in order to deposit an egg pod ca. 8-10 cm deep into the ground (Fig. 1A). This is aided by two pairs of dedicated sclerotized digging valves, or oviposition valves (Fig. 1B), which are utilized for excavating a deep and narrow burrow (Vincent, 1976; Das et al., 2022a). After completing the digging, the female begins to secret into the bottom of the burrow a small amount of a specialized protein-based foam, followed by depositing the egg pod. Subsequently, while concomitantly withdrawing her abdomen, she fills the burrow to the top with this specialized foam, which is intended (in addition to its other roles, e.g. Lavy et al., 2021) to provide the future hatchlings with a quick and safe route to the surface (Symmons and Cressman, 2001; Uvarov, 1977; Hägele et al., 2000). The female locust abdomen at rest is no longer than 2.5-3 cm. During oviposition digging, however, it demonstrates extreme extension, 2-3 fold its original length (Fig 1; Vincent, 1975; Jorgensen and Rice, 1983; Das et al., 2022b). The locust achieves this by means of a remarkable extension of the soft tissue within the abdomen. This includes extension of parts of the intersegmental membranes up to 10 fold their original length (Vincent and Wood, 1972; Vincent, 1975) and a similar extreme elongation of certain abdominal intersegmental muscles (Jorgensen and Rice, 1983). This elongation of the female locust’s abdomen is accompanied by an extraordinary longitudinal stress-induced elongation of the abdominal nervous system (Das et al., 2022b).

**Figure 1.**
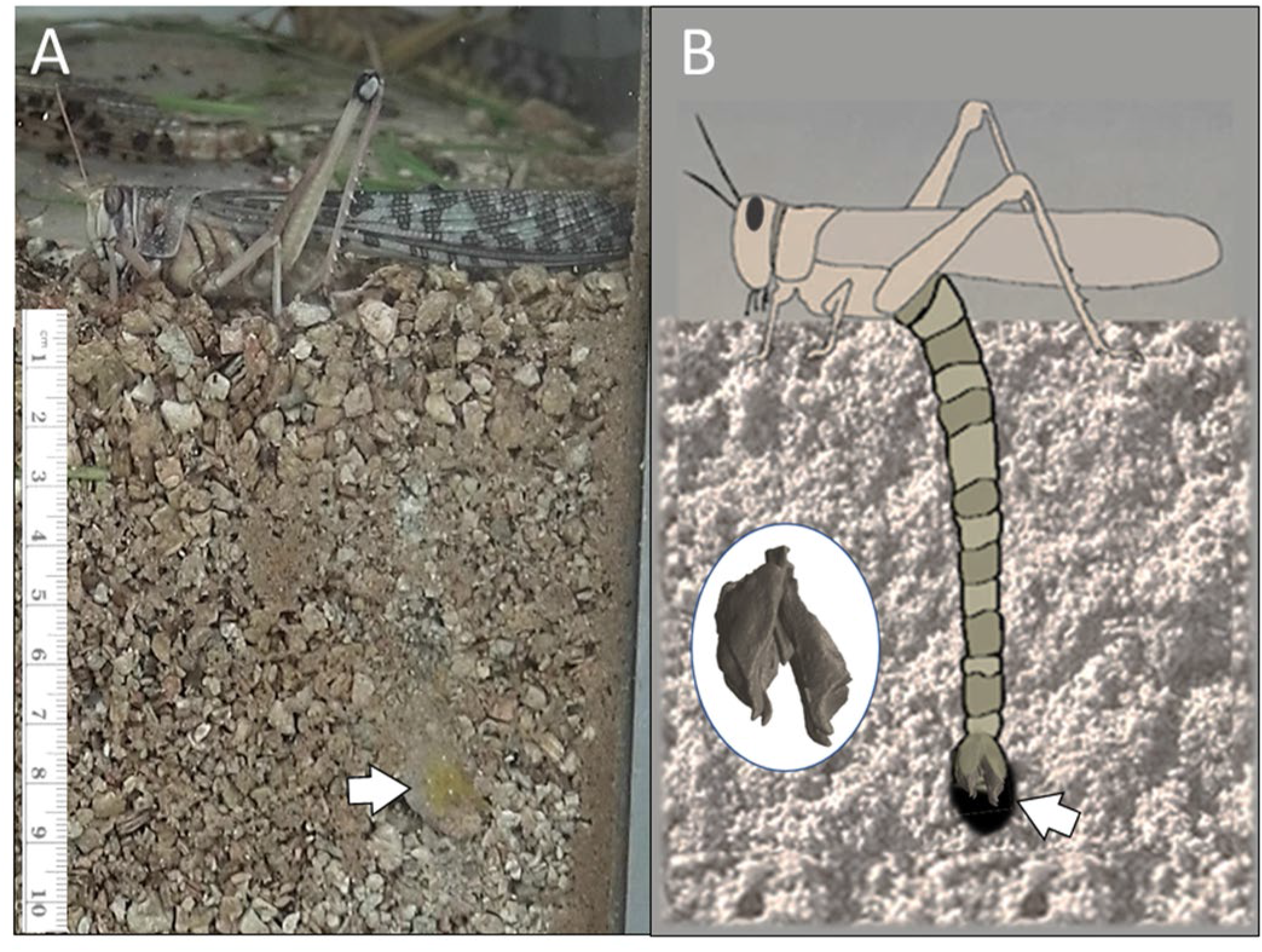
**A**. A snapshot from a video sequence depicting oviposition digging in the female locust (taken through a glass wall). The locust is shown after completing the burrow, as she starts retracting her abdomen, discharging the foam secretion and depositing the eggs (arrow). B. A schematic illustration of the female’s abdomen during excavation, depicting the digging apparatus – the ovipositor valves and the extreme extension of the abdomen.

Both laboratory studies and field observations have noted that gravid locust females are fastidious when choosing a site at which to deposit their eggs (Kennedy, 1949; Popov 1958, Hunter-Jones 1964, Pener & Simpson 2009, Tanaka & Sugahara 2017), often delaying oviposition for up to 72 hours. If a suitable site is not found within this period, the locusts will dispose of their eggs in the open, avoiding the costly procedure of digging an unsuitable site and its associated dangers of predation and being trapped (if the female cannot withdraw her abdomen after digging during oviposition). Almost all previous work, however, has focused on the suitability of the oviposition site in relation to the presence of predators, shade, temperature, and, mostly, humidity (all critical factors for the survival of the female, her eggs, and hatchlings). In contrast, there is very little knowledge regarding the ability of the female to negotiate any physical obstacles encountered within the substrate during oviposition digging.

Here we sought tot answer the question of how the active digging oviposition valves negotiate obstacles encountered in their path. To investigate this, we provided gravid females in our gregarious locust breeding colony with access to PVC oviposition tubes filled with moist sand (Fig. 2). Some of these tubes were prepared with specialized 3-D printed plastic obstacles (mimicking natural obstacles like roots and pebbles). The females’ digging paths were then reconstructed and analyzed. Our findings revealed a remarkable ability of the locust to circumvent such impediments by steering its digging valves and abdomen around the obstacles and maintaining a general vertical digging path.

**Figure 2.**
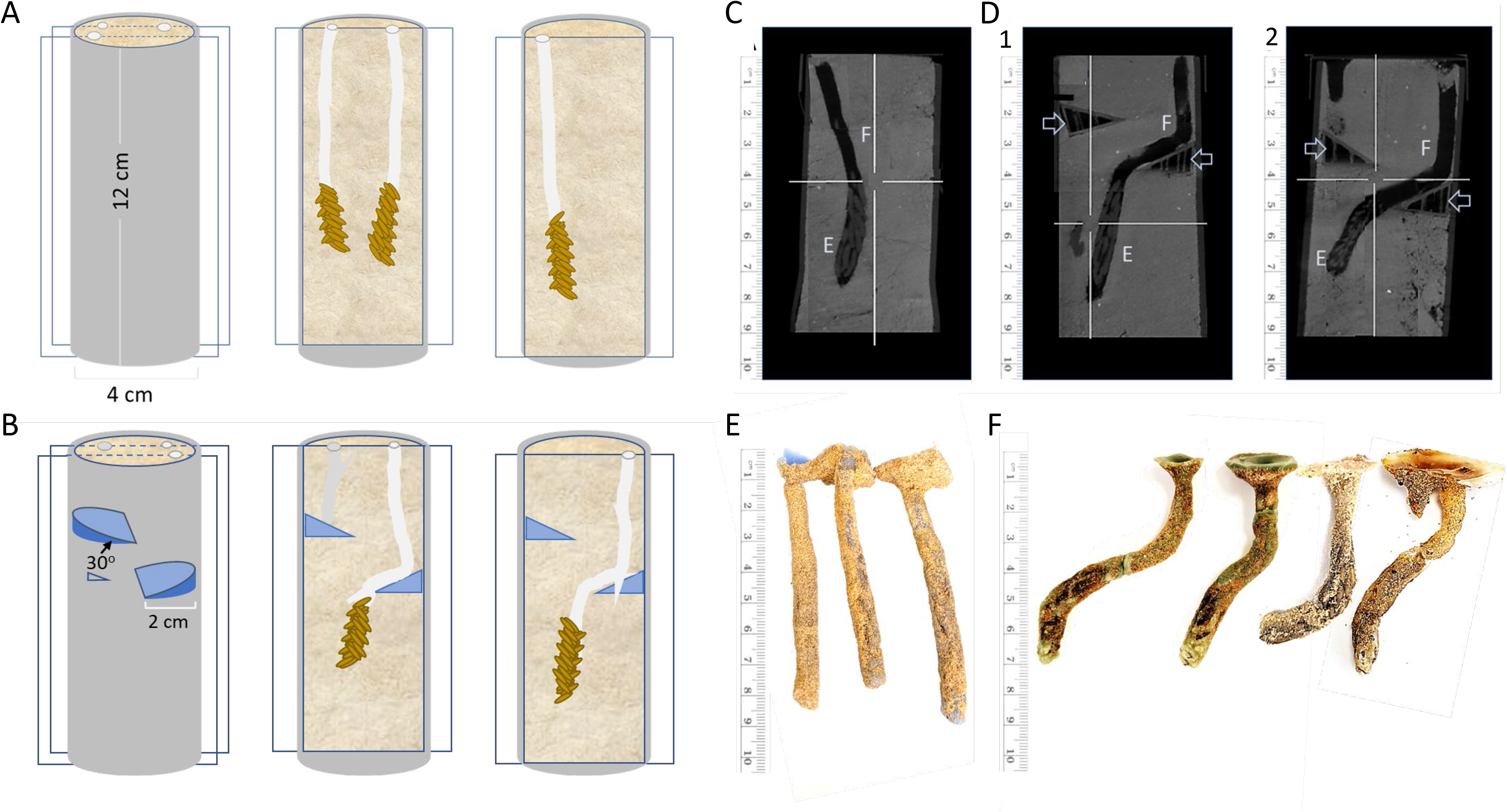
The oviposition digging path. **A**. A schematic illustration depicting the oviposition tubes and two sagittal-longitudinal sections drawn to demonstrate examples of digging paths, including egg pods (orange) and foam secretion (white), in control conditions. **B**. A similar schematic illustration of an experimental oviposition tube (the two plastic obstacles are shown in blue), and sagittal sections depicting two examples of the oviposition digging when the female encountered an obstacle. One failed attempt is also shown (left on the left section). **C-D** Monitoring and analyzing the oviposition digging path. Composite images, i.e., overlays of several view plains, from examples of HDCBCT scans of sand-filled PVC oviposition tubes: Control **(C**; Foam plug and egg pod are marked by F and E, respectively) and tubes prepared with 2 plastic obstacles (arrow heads) at 2 and 4 cm (**D1**) or at 3 and 5 cm (**D2**). Foam plug and egg pod are noted with a F and E respectively. **E** and **F**. Paraffin casts of the digging path remaining after the eggs hatched and the hatchlings surfaced via the foam plug. The relatively straight control digging (**E**) is compared to the obstacle circumvention in the experiments (**F**).

## Materials and Methods

Desert locusts, *Schistocerca gregaria* (Forskål), in our breeding colony at the School of Zoology, Tel Aviv University, are reared for many consecutive generations under heavy crowding, 100-160 animals in 60-liter metal cages. The cages are kept under controlled temperature and humidity conditions (30°C, 40-60%) under a 12D:12L cycle. Direct radiant heat is supplied during daytime by incandescent electric bulbs, to reach a final day temperature of 35-37°C. The locusts are fed daily with wheat seedlings and dry oats.

Cages with gravid females were supplied with 12×4cm PVC or cardboard tubes densely filled with wet sand for oviposition. The tubes were either control or experimental tubes (Fig. 2). The latter contained two specially constructed and positioned 3D-printed plastic obstacles (Fig. 2B), at depts of either 2 and 4 cm, or 3 and 5 cm. The oviposition tubes were left overnight and checked the next day for signs of oviposition: holes in the sand indicating unsuccessful attempts, and traces of the secreted foam (foam plug) indicating completed oviposition events (Figs. 2).

The females’ oviposition digging paths were then studied and characterized in one of three approached:

1. *Visual inspection*. The cardboard tubes were carefully cut longitudinally, and the sand was gently scraped with a scalpel to expose the foam and egg pod along the digging path. Various aspects of the digging path geometry were documented, including length (depth), changes in the path direction, and deviation angles from a vertical-linear line.
2. *Cone Beam Computed Tomography (CBCT) scans in high definition (HD) mode*. This was carried out at the Tel Aviv University Faculty of Medicine, School of Dental Medicine. CBCT scans were performed with an Ortophos 3D SL (Sirona Systems GmbH, Bensheim, Germany) in HD mode using the following protocol: field of view - 11 cm x 10 cm, X-ray tube voltage −85 kV, X-ray tube current - 7 mA, isotropic voxel edge length - 160 μm, effective radiation time - 4.4 s. The resulting images were analyzed using native Sidexis 4 (v. 4.3.1, Sirona Systems GmbH, Bensheim, Germany) software.
3. *Paraffin casts*. The oviposition tubes were incubated (11 days at 37°C) and the eggs were allowed to hatch. The hatchlings were collected at the surface, leaving behind them the open hole and the burrow previously plugged with the foam that the hatchlings use as their route to the surface. Ordinary, commercially available paraffin wax was heated to a liquid state and carefully poured onto the surface of the sand tubes, allowing it to penetrate and completely fill the holes, all the way down to the hatched eggs. The paraffin was then allowed to harden and solidify before it was carefully extracted from the surrounding sand.

## Results

We visually inspected ca. 45 oviposition tubes, each comprising 1-5 cases of successful oviposition digging. Visual inspection data was augmented by the paraffin casts (overall 15 oviposition tubes) and by our HD-CBCT imaging (overall 8 oviposition tubes).

During oviposition digging the female locusts demonstrate positive geotropism, i.e., they tend to dig and burrow in a path that is sometimes slightly curving, but mostly close to straight-vertical (Figs 1). Accordingly, digging in the control tubes (no obstacles), was characterized by an initial practically vertical (4.0° ±5.7° deviation from vertical) and a nearly straight digging path (final path showing 2.7° ±4.2° deviation from the initial angle), to an average depth of 9.4±1.1 cm (Figs 2A, C, and E, n=15).

We next challenged the locusts with tubes that contained obstacles comprising a 30° inclined upper surface. Our examination of the experimental oviposition tubes that contained an obstacle at 2 cm depth, revealed that upon encountering such an obstacle, the females mostly abandoned digging and withdrew their abdomen. Such failed attempts were evident only by the relatively shallow hole in the sand. In contrast, successful oviposition was often observed in tubes that contained obstacles at 3-5 cm depth. In marked contrast to the control, in many such cases the digging path showed bends and turns: first, to bypass the obstacle; and, in some cases, a second turn to restore an overall vertical inclination (Figs. 2B, D, and F). This complex path was prevalent upon encountering an obstacle at 3 or 4 cm depth, but was also seen for some of the 5 cm deep obstacles. Overall, we have found a clearly curved digging path in 18 cases, out of which 6 showed a second turn restoring the vertical path. The initial observed digging path showed a 15.1° ±10.5° deviation from vertical, followed by a first turn of 33.0° ±6.6° and (when present) a second, counterturn of 38° ±5.9°. In all cases, the diverted digging path was adjacent to the obstacle slope. The average length of the digging path was found to be 9.2±2.7.

In order to unravel the dynamics of the obstacle negotiation by the locust females, we conducted another set of experiments, in which the upper surface of the obstacles (3 and 5 cm deep) was horizontal (perpendicular to the initial digging path), and not angled at 30° as in the initial set of experiments. In these latter experiments (n=13), we observed further evidence of excavation attempts that terminated upon reaching the obstacle (failing to circumvent it; Fig. 3), or after “miscalculating” and turning towards the walls of the tube rather than to the center. Even clear cases of horizontal digging were observed (Fig. 3D). Some experiments, however, also presented examples of successful oviposition digging (n=6). These revealed an S-shaped winding path, avoiding both the first and second obstacles (Fig. 3). Interestingly, unlike the cases of the sloped-surface obstacles, here the digging path did not touch or flank the obstacles, suggesting that the female had performed some retraction and rerouting of her digging upon encountering the obstacles. The complex nature of these digging paths, however, challenges their representation in 2D (Fig. 3).

**Figure 3.**
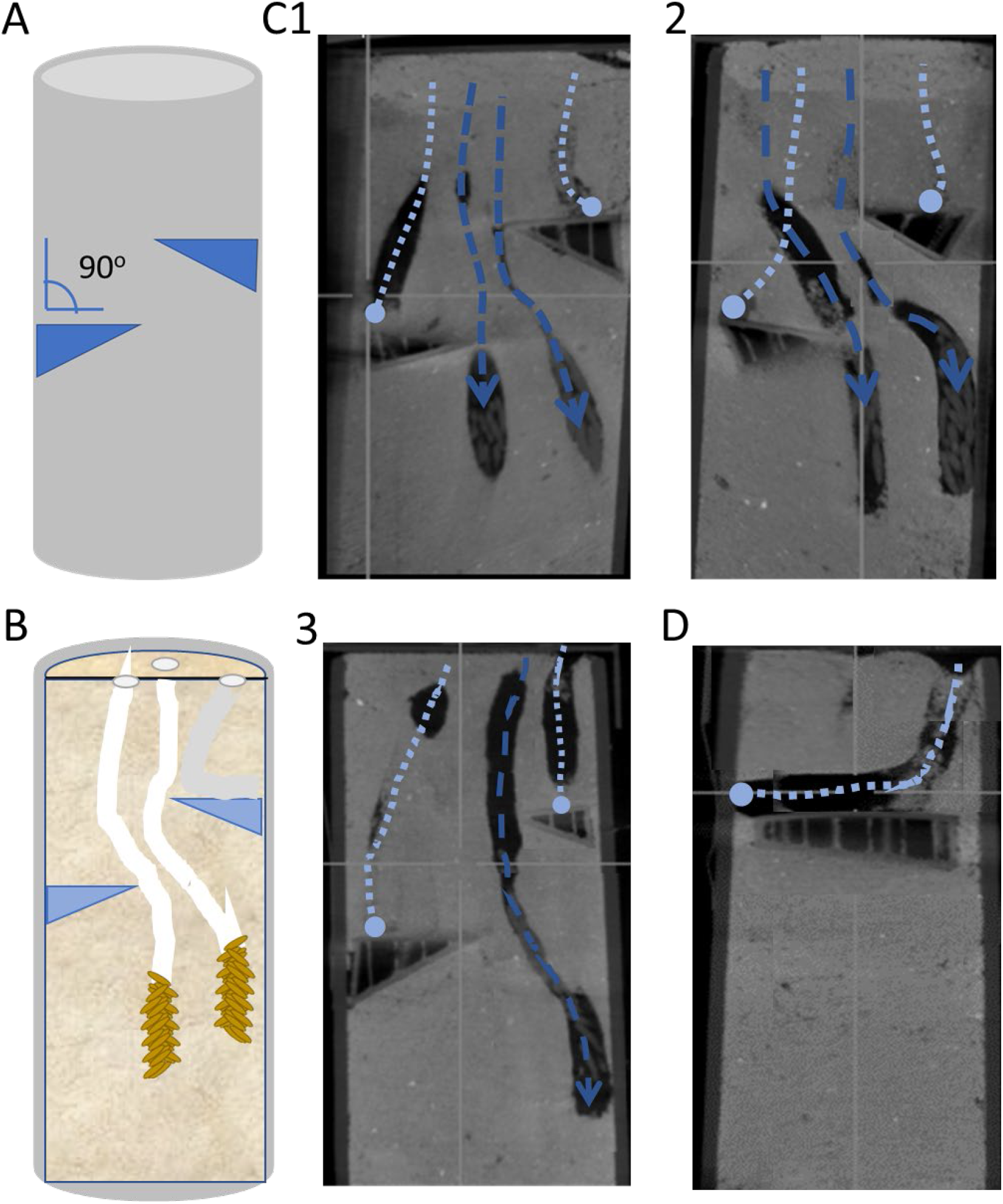
Experiments involving obstacles presenting an upper surface perpendicular to the initial vertical path. **A**. A schematic illustration of the experimental oviposition tube. **B**. A sagittal-longitudinal section drawn to demonstrate examples of digging paths, including egg pods (orange) and foam secretion (white). **C1-3**. Optical sections from HD CBCT scans of three experiments showing failed oviposition attempts (light blue dotted lines), and digging paths successfully avoiding the obstacles (blue, dashed lines; arrow head depicting the egg pod). **D**. An exceptional example of a failed oviposition attempt involving extended horizontal digging.

## Discussion

In this study we challenged female locusts with two types of obstacles in order to examine and characterize their ability to negotiate subterranean obstacles during oviposition digging: one obstacle presented a sloped surface (30°), and the other presented a horizontal surface - perpendicular to the initial digging path. When encountering the former, the females demonstrated a digging path that closely followed the obstacle surface. Subsequently, having cleared it, they resumed their vertical inclination, suggesting that these changes in digging direction involved some sort of sensory-motor decision-making and did not merely constitute physical interaction with the obstacle. Introduction of the horizontal-surface obstacles somewhat increased the probability of failed digging attempts, i.e. subsequent retraction of the abdomen and abandoning the dig. Nevertheless, successful cases of oviposition digging and egg deposition were also demonstrated, in which the females clearly made “the right” choice regarding the direction to take in order to circumvent the obstacle. The reorientation of the digging was mostly preceded by some retraction of the abdomen, resulting in an intricate digging path that was not closely adjacent to the obstacles. Making a wrong choice (leading to excavation in the direction and along the walls of the PVC tube) resulted in the female demonstrating its remarkable ability for extreme bending and digging in a horizontal orientation.

In her exhaustive and extremely thorough series of reports on oviposition digging in the grasshopper, including its anatomy, the motor program, and neural control, Thompson (1986a,b) notes that: “*Once the ovipositor has become engaged in the substrate, the female simply stands on the surface while the ovipositor burrows beneath her*”. This is a quite accurate description, although it downplays the amazing feat that is demonstrated by the insect, including the use of a unique digging mechanism, as well as the far yet from fully explained extreme elongation and extension of the abdomen. Most importantly, it assumes that the substrate is simple and homogenous, neglecting the sometimes intricate steering and obstacle negotiation, which is very often required in natural settings.

Overall, bending movements of the locust’s abdomen are controlled by the intersegmental longitudinal muscles (Snodgrass, 1935). Notably, these longitudinal muscles are largely suppressed during oviposition digging (Rose etal., 2000). This is in order to avoid resistance to the extreme extension of the abdomen (Vincent, 1975; Das et al., 2022b), to restrict the forces working against the increased internal pressure in the abdominal segments and, mostly, to eliminate any resistance to the pulling forces exerted by the oviposition valves as they anchor the abdominal tip in the substrate (Jorgensen and Rice, 1983). Hence, the mechanism behind the ability of the female locust’s abdomen to steer free of obstacles encountered in a non-homogenous substrate during oviposition digging, is probably mostly based on the direct musculature of the ovipositor valves.

Locusts and grasshoppers are unique in possessing ovipositor valves derived from appendages and are the only reported insects whose ovipositor works by opening and closing movements rather than by sliding valves upon each other (Snodgrass, 1935; Matsuda, 1976). Snodgrass (1935) showed that the two pairs of digging valves are hinged at their bases to each other and to a prominent pair of internal apodemes. The valves’ movements were reported to involve the combined action of ten pairs of muscles in abdominal segments eight and nine (Nel, 1929). These, in turn, are controlled by 17 ovipositor motor neurons that, together with a few additional muscles in other abdominal segments, and a digging central pattern generating circuit (CPG) located in the VII^th^ and VIII^th^ abdominal ganglia, are all involved in the complex cyclical rhythm of protraction, opening, closing, and retraction of the ovipositor valves (Ayali & Lange, 2010). In practically all studied CPGs, the motor pattern is modulated by sensory inputs (e.g., Ayali et al., 2015, and references within). One prominent such input, in the case of the digging CPG, is provided by the mechanosensory hairs covering the ovipositor valves (Belanger and Orchard 1992). Kalogianni (1995) reported that the hairs on the external surfaces of the ventral and dorsal ovipositor valves respond to wind stimulation, whereas the hairs on the inner surfaces of the dorsal valves are not wind-sensitive. All ovipositor hairs, however, respond to tactile displacement (Kalogianni, 1995), and are thus well suited to detect obstacles in the substrate. Such sensory input, in turn, can be instrumental in inducing lateral movements that will redirect the overall digging in a new direction, bypassing an obstacle.

There is also the possibility that the steering of the abdomen tip may be an indirect passive outcome of the movement dynamics and forces exerted by the valves: i.e., that the above-noted cycle of movements of the valves is sufficient to steer the abdomen sideways upon working against an obstacle, avoiding the need for an intricate sensory-motor integration process. In other words, the only type of (pseudo) decision-making involved, is an embodied one, i.e., one that results directly from the digging valves’ morphology and movement dynamics. While such a mechanism may indeed be involved in obstacle negotiation during oviposition, it seems insufficient to explain our findings, being mostly not supported by the results of the experiments in which we observed two bends or turns in the digging path: a first one that follows the surface of the obstacle; but then also a second one that restores an overall vertical inclination. Our findings in the experiments in which the obstacles’ surface was perpendicular to the digging path are clearly more in accord with a process that involves decision-making and central control.

In summary, obstacle negotiation appears to be a crucial aspect of the subterranean oviposition behavior in female locusts. Our findings reveal that the oviposition process is significantly more complex than previously understood and described. This complexity involves sophisticated sensory-motor integration, encompassing obstacle sensing and directional steering, probably under meticulous central control.

## Acknowledgments

We are grateful to Bat-El Pinchasik and Shai Sonnenreich for the constructive discussions. We thank Ben Maoz for the valuable assistance in 3D printing; and Rakesh Das and Omer Lavy for contributing to the illustration in Figure 1B.

## Notes

### Competing Interest Statement

The authors have declared no competing interest.

